# Targeting IDH1-Mutated Oligodendroglioma with Acid Ceramidase Inhibitors

**DOI:** 10.1101/2024.04.27.591426

**Authors:** Helena Muley, Tyrone Dowdy, Faris Zaibaq, George Karadimov, Aiguo Li, Hua Song, Meili Zhang, Wei Zhang, Zalman Wong, Laurence Zhang, Adrian Lita, Mioara Larion

## Abstract

Oligodendroglioma is genetically defined as a tumor harboring isocitrate dehydrogenase 1 or 2 mutations (IDH1^mut^/IDH2^mut^) and 1p/19q co-deletions. Previously, we reported that in IDH1^mut^ gliomas, D-2HG, the product of IDH1 mutant enzyme produces an increase in monounsaturated fatty acid levels that are incorporated into ceramides, tilting the S1P-to-ceramide rheostat toward apoptosis. Herein, we exploited this imbalance to further induce and IDH^mut^-specific glioma cell death. We report for the first time that the inhibition of acid ceramidase (AC) induces apoptosis and provides a benefit in mice survival in IDH1^mut^ oligodendroglioma. We demonstrated an IDH1^mut^-specific cytotoxicity of SABRAC, an irreversible inhibitor of AC, in patient-derived oligodendroglioma cells. Exploring the mechanism of action of this drug, we found that SABRAC activates both extrinsic and intrinsic apoptosis in an ER stress-independent manner, pointing to a direct action of AC-related ceramides in mitochondria permeability. The activation of apoptosis detected under SABRAC treatment was associated with up to 30-fold increase in some ceramide levels and its derivatives from the salvage pathway. We propose that this novel enzyme, AC, has the potential to increase survival in oligodendroglioma with IDH1^mut^ and should be considered in the future.

## Introduction

Oligodendroglioma, although rare, is the third most common glioma in adults ^1^. An estimated 1,217 people with oligodendroglial tumors are diagnosed every year in the United States, most often in people between the ages of 35 and 44. Diagnosis of oligodendroglioma is based on the finding of two genetic alterations: mutations in one of two genes called isocitrate dehydrogenase 1 and 2 (IDH1^mut^/IDH2^mut^) and the absence of the two chromosomal arms, 1p and 19q ^2^. Although patients with IDH^mut^ glioma exhibit a longer survival than the ones with IDH-wild-type (IDH^wt^) glioma, this brain tumor still recurs and may undergo malignant transformation to a higher grade. Additionally, oligodendrogliomas are more prone to develop a hypermutation phenotype which is also associated with a worse prognosis ^3^. The current therapies for even the highly treatment-responsive oligodendrogliomas are not curative ^4,5^, highlighting that there is an urgent need for more effective and personalized treatments. While many studies are evaluating a variety of cytotoxic, molecular, and immunologic therapies, few are examining potential unique metabolic vulnerabilities in oligodendroglioma.

Sphingolipids are one of the major lipid families in human cells, with a wide range of functions in the organization of cell membranes and signal transduction ^6^. Most of the investigation into sphingolipid signaling has focused on the dynamic balance and opposing nature of intracellular ceramide and sphingosine-1-phosphate (S1P), also called “sphingolipid rheostat”. Ceramide is associated with growth arrest and apoptosis, while S1P promotes cell proliferation and survival ^7,8^. The rheostat S1P-to ceramide has been well described in glioblastoma (IDH1^wt^) and is associated with many of its hallmarks ^9^. In fact, lower ceramide and higher S1P levels correlate with higher glioma grades ^10^. However, the role of this rheostat in IDH1^mut^ gliomas has not been widely investigated.

IDH1 wild-type enzyme catalyzes the conversion of isocitrate to α-ketoglutarate (αKG), whereas the mutant IDH1 protein exhibits neomorphic activity, converting αKG into D-2-hydroxyglutarate (D-2HG). Previously, we demonstrated that in IDH1^mut^ glioma, D-2HG, the product of IDH1 mutant enzyme produces an increase in monounsaturated fatty acid (MUFA) levels ^11^. These increased MUFA are incorporated into ceramides, tilting the S1P-to-ceramide rheostat toward apoptosis ^12,13^. Therefore, for the first time, it has been described the rheostat imbalance in IDH^mut^ gliomas, to be the opposite of that known in IDH^wt^ gliomas. Moreover, this imbalance can be manipulated by decreasing S1P or increasing ceramides to the point that leads to IDH^mut^-specific cell death ^12,13^.

To exploit this metabolic vulnerability, we inhibited AC, an enzyme in the sphingolipid pathway, and after testing several inhibitors, we identified SABRAC as one of the top drugs inducing cytotoxicity in IDH^mut^ glioma cells. SABRAC is a specific and irreversible inhibitor of AC ^14^. Ceramidases are key enzymes that maintain the intracellular homeostasis of ceramide decreasing their levels ^15^. Herein, we report that IDH mutant gliomas, and especially oligodendroglioma have a lower expression of AC compared to IDH wild-type gliomas. This reduced expression is a vulnerability of this type of tumors making them more sensitive to AC inhibitors. We demonstrated how AC inhibitors as SABRAC, increase ceramides in oligodendrogliomas cells leading them to mitochondrial-driven apoptosis. Interestingly, the induced sphingolipid pathway was accompanied by a sharp rise in cholesterol. Consistently, changes in the expression of genes related to ceramide and cholesterol signaling and pathways were found impaired. Moreover, treating with SABRAC mice that harbor IDH1^mut^ oligodendroglioma intracranially, significantly increases their survival. Our study thus reveals an IDH mutant-specific toxicity of SABRAC and suggests that inhibiting AC may represent a promising therapeutic strategy for treating oligodendroglioma.

## Methods

### Cell models

The human glioma cells used in this project were: TS603 (grade III oligodendroglioma, kindly provided by Dr. Timothy Chan’s lab), NCH612 (grade III oligodendroglioma, Cytion), BT142 (grade III oligoastrocytoma, ACS-1018 ATCC), GSC923 (grade IV glioblastoma, NOB lab), and U251MG (grade IV glioblastoma, ATCC). U251^*WT*^ and U251^*R132H*^ (cells were previously produced by Dr. Yang in NOB). U251 cells were cultured in DMEM (10-013-CV Corning) supplemented with 10% fetal bovine serum (SH12450H R&D Systems) and 1% penicillin-streptomycin (15140122 Gibco). Non-immortalized human astrocytes were obtained from ATCC.

### Organoid culture

All the experiments were performed using human-derived stem-like spheroids. Neurospheres were grown in suspension in DMEM/F12 medium (11320033 Gibco) supplemented with 1% N2 growth supplement (17502048 Gibco), 2μg/ml heparin sodium salt (07980 Stem Cell), 20ng/ml EGF (236-EG R&D Systems), 20ng/ml FGF (3718-FB R&D Systems) and 1% penicillin-streptomycin (15140122 Gibco).

### Cell viability assays

For cell viability assays, cells were plated using optimal seeding densities per well in 96-well plates with neurosphere growth medium and drug treatments. After 48 hours, cell viability was assessed by Cell Counting Kit-8 (CK04 Dojindo). The absorbance was measured at 450 nm. The concentration of the drug resulting in 50% inhibition of cell viability (IC50) was calculated using non-linear regression curve fitting.

### Immunoblotting

Cell lysates were prepared using NP-40 lysis buffer (J60766.AP Thermo Scientific) containing protease inhibitors (A32955 Thermo Scientific). The lysates were centrifuged at 13,000 × g for 10 min to obtain the supernatants. BCA assays (23225 Thermo Scientific) were performed for measuring protein concentration. The proteins in the lysates were resolved by SDS-polyacrylamide gel electrophoresis and were transferred to LF PVDF membranes. The immobilized proteins were immunoblotted with antibodies against the proteins of interest. The bands were visualized using an enhanced chemoluminiscence (ECL) system (ChemiDoc MP BIO-RAD). Equal loading of proteins was verified by immunoblotting for β-actin or α-tubulin. All the antibodies used in immunoblot analysis are listed below. Rabbit anti-ASAH1 (HPA005468 Sigma), rabbit anti-Caspase 8 (13423-1-AP Proteintech), rabbit anti-BCL-2 (12789-1-AP Proteintech), rabbit anti-BAX (5023 Cell Signaling), rabbit anti-Cytochrome c (11940T Cell Signaling), rabbit anti-Caspase 3 (19677-1-AP Proteintech), rabbit anti-cleaved PARP1 (ab32064 abcam), rabbit anti-phospho-BAD (9291S Cell Signaling), rabbit anti-BAD (ab32445 abcam), rabbit anti-p8 c-terminal (SAB2109172 Sigma), rabbit anti-phospho-eIF2α (9721S Cell Signaling), rabbit anti-eIF2α (9722S Cell Signaling), rabbit anti-PERK (3192S Cell Signaling), rabbit anti-CHOP (15204-1-AP Proteintech), rabbit anti-ATF4 (11815S Cell Signaling), rabbit anti-TRIB3 (13300-1-AP Proteintech), mouse anti-beta actin (ab6276 abcam), peroxidase-conjugated AffiniPure goat anti-rabbit IgG (111-035-144 Jackson) and peroxidase affiniPure goat anti-mouse IgG (115-035-146 Jackson).

### LC/MS sample preparation

TS603 oligodendroglioma cells (5 × 10^6^ cells per condition) were treated with DMSO, 5μM, 10μM or 20 μM of SABRAC for 48h. Cells were collected by centrifugation at 300 x g for 5 min and washed with PBS. All remaining PBS was removed using an additional centrifugation step.

### Sphingolipid and polar lipid optimized extraction

Samples extraction was performed as described in Dowdy T. et al, 2020 ^12^. The only modification from the protocol was related to the solvent composition used for reconstituting the hydrophilic and hydrophobic phases that was 100 μL of 5:4:1 EtOH/MeOH/water.

### LC/MS analysis

LC/MS analysis were previously described in Dowdy T. et al, 2020 ^12^, except for the mass accuracy and retention time values. Targeted ion selection, alignment, and annotation for logical binning of the input data were restricted to ion mass accuracy ± 2.0 mDa and retention time ± 0.5 min using an in-house Personal Compound Data Library (PCDL) of known sphingolipids. Following pre-processing, the ion area for each analyte was adjusted to ratio by sample-specific internal standard area, and then corrected to sample-specific weight quantification.

### RNA extraction

TS603 oligodendroglioma cells (2.5 × 10^6^ cells per condition) were treated with DMSO, 2.5μM or 5μM of SABRAC for 48h. We checked by western blot that the cells were not in apoptotic conditions. Total RNA was purified using the PureLink™ RNA Mini Kit (12183025 Invitrogen). An additional step of DNA digestion was added using the PureLink™ DNase Set (12185010 Invitrogen). The integrity of the RNA was assessed using a 5300 Fragment Analyzer System (Agilent Technologies). RNA samples with RNA Integrity Number greater than 9.7 were submitted to the CCR Sequencing facility (https://ostr.ccr.cancer.gov/resources/sequencing-facility/). Paired-end *sequencing* (2x100bp) was performed on a *NovaSeq* 6000 instrument using the *NovaSeq* 6000 *S1* Reagent Kit.

### RNA-seq data analysis

Sequencing reads of RNA-seq raw data were analyzed using CCBR Pipeliner (https://github.com/CCBR/Pipeliner). The pipeline performs several tasks: pre-alignment reads quality control, grooming of sequencing reads, alignment to human reference genome (hg38), post-alignment reads quality control, feature quantification, and differentially-expressed gene (DEG) identification. In the QC phase, the sequencing quality of each sample is independently assessed using FastQC, Preseq, Picard tools, RSeQC, SAMtools and QualiMap. Sequencing reads that passed quality control thresholds were trimmed for adaptor sequences using Cutadapt algorithm. The transcripts were annotated and quantified using STAR. Sample quality control was vigorously carried out based on sequencing criteria such as mapping rates, library complexities, etc. Further sample filtering was performed based on zero value of specific genes and the sample variance. DEGs were determined using DESeq2 in idep (http://bioinformatics.sdstate.edu/idep96/) with the thresholds of p < 0.05 and |FC| =1.3.

### Functional Pathway and Network Analysis

Pathway and network analysis was carried out using Ingenuity Pathway Analysis (IPA; www.qiagenbioinformatics.com). The DEGs (either up-or downregulated) were uploaded into IPA and core analysis was performed for each comparison. Enriched canonical pathways and network analysis were inspected to elucidate underlying molecular functions.

### Bioinformatic analysis of datasets of human glioma samples

Gene expression and survival of adult glioblastoma patient data from the TCGA Project (TCGA-GBMLGG) and the Chinese Glioma Genome Atlas (CGGA, http://www.cgga.org.cn/) ^16^, were analyzed using GlioVis data portal (http://gliovis.bioinfo.cnio.es/) ^17^.

### In vivo oligodendroglioma xenograft model

For SABRAC toxicity study, 16 eight-week-old female SCID mice were randomly divided into four groups having one vehicle control group (n=4) and three SABRAC-treatment groups (n=4 for each dose; 1mg/kg, 5mg/kg and 15mg/kg). Each animal received five intraperitoneally injections per week of SABRAC (SABRAC in 0.5% methylcellulose and 0.2% Tween 80 in PBS) or vehicle control (DMSO in 0.5% methylcellulose and 0.2% Tween 80 in PBS). Routine cage-side observations were made on all animals at least once a day throughout the study for general signs of pharmacologic and toxicologic effects, morbidity, and mortality. Mice were weighed every 3-4 days. 4h after the last injection mice were euthanized and plasma, liver and brain were collected. Brains were processed and analyzed by LC/MS.

For evaluating the efficacy of SABRAC in the oligodendroglioma xenograft mice model, 200,000 human TS603 oligodendroglioma cells were intracranially injected in 18 eight-week-old female SCID mice. One week after the injection, mice were randomly divided into two groups having a vehicle control group (n=9) and a SABRAC-treatment group (n=9, 15mg/kg). Each animal received five intraperitoneally injections per week of SABRAC (SABRAC in 0.5% methylcellulose and 0.2% Tween 80 in PBS) or vehicle control (DMSO in 0.5% methylcellulose and 0.2% Tween 80 in PBS). Routine cage-side observations were made on all animals at least once a day throughout the study for general signs of pharmacologic and toxicologic effects, morbidity, and mortality. Mice were weighed every 3-4 days. Animals were euthanized when reaching the endpoint of deteriorated clinical score and weight-loss (more than 15% of body weight).

### Statistical analysis

Statistical significance was determined using the appropriate test according to the samples and study design. Prior to the comparisons, the normality of the distributions was tested with the Shapiro-Wilk test before comparison. All data were presented as the mean ± standard error or the mean ± SEM of at least three independent experiments unless otherwise indicated in the figure legend. p values less than 0.05 were considered statistically significant. Sample size and p-values are indicated in main and supplementary figure legends. Statistical analysis for human glioma samples was obtained from GlioVis ^17^. Log-rank and Wilcoxon tests were applied for Kaplan–Meier survival curves, and Tukey’s Honest Significant Difference (HSD) was applied to compare ASAH1 expression across tumor histology. For LC-MS data, statistical analysis was performed using Metaboanalyst 6.0 software ^18^. The corrected values for each analyte were log-transformed and normalized to the median to perform statistical analysis using non-parametric, Mann-Whitney U t-test for binary comparisons. For RNA sequencing analysis, DEGs were determined using DESeq2 in idep with the thresholds of p < 0.05 and |FC| =1.3. The rest of statistical analyses were performed with GraphPad Prism software (version 10.0). Friedman and Kruskal-Wallis tests were applied. For mice experiments, Log-rank (Mantel-Cox) statistical test was applied for Kaplan-Meier survival analysis and two-way ANOVA for body weight analyses.

### Graphical abstracts

Illustrations were performed using BioRender.com.

## Results

### High expression of ASAH1 is associated with poor survival in patients with oligodendroglioma

In IDH1^mut^ gliomas, the S1P-to-ceramide rheostat is tilted towards high ceramide levels (Fig. 1A, ^12,13^). Taking advantage of this vulnerability, we inhibited acid ceramidase (AC), with specific inhibitors that were expected to increase ceramides. As a result, we identified SABRAC, a specific and irreversible AC inhibitor ^14^, as one of the top drugs that induced cytotoxicity in IDH^mut^ glioma cells (Fig 1A, Fig S1A ^12^). AC hydrolyzes lysosomal membrane ceramide into sphingosine, the backbone of all sphingolipids ^19^, (Fig. 1B). Next, we decided to explore the expression of AC in different brain tumors. Using The Cancer Genome Atlas (TCGA) data, we found that AC expression is significantly lower in IDH^mut^ gliomas, especially in oligodendroglioma, compared to IDH^wt^ gliomas (Fig. 1C). We also found that high AC expression is associated with worse survival only in oligodendroglioma (Fig. 1D). To see if IDH1 mutation was modulating AC expression, we performed western blots of IDH wild-type and mutant glioma cells. Interestingly, the presence of the mutation was related to decreased AC expression. Moreover, a U251 glioblastoma model overexpressing IDH1 mutation (R132H), presented reduced levels of AC compared to the empty vector (Fig. 1E), suggesting that IDH1^mut^ decreases AC levels. Since five ceramidases have been identified in humans, we explored the expression of all five ceramidases in oligodendroglioma tumors in TCGA data, and we identified AC as the ceramidase with higher expression. Next, we compared the mRNA levels of all five ceramidases in TS603 PDX-derived oligodendroglioma cell line that we profiled using RNA sequencing, and we found similar results to TCGA, where AC, had the highest mRNA levels amongst all five enzymes (Fig. 1F).

**Fig. 1.**
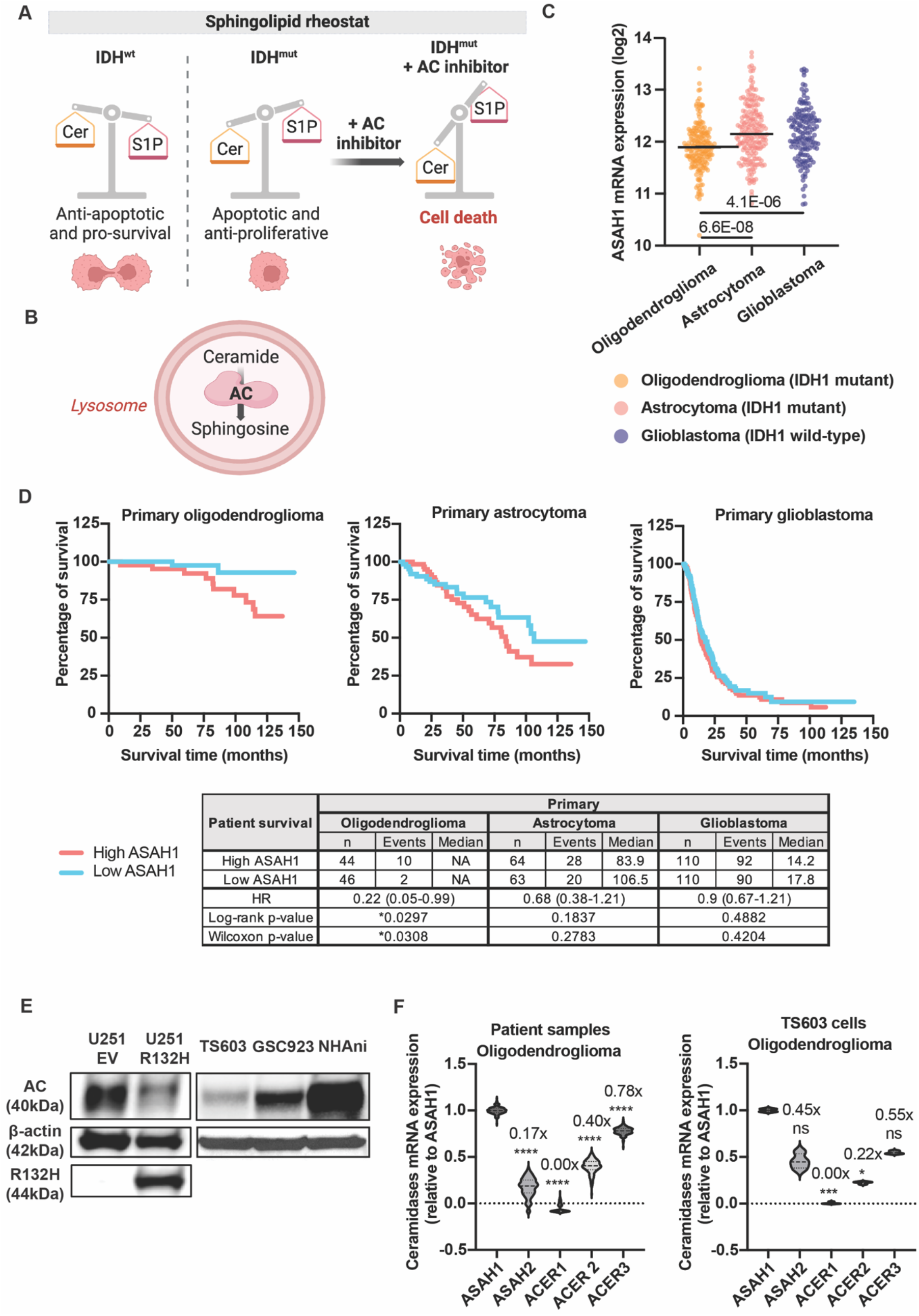
The lower expression of ASAH1 in IDH-mutant compared to IDH-wild-type gliomas, is associated with a better survival. A. Image illustrating how the sphingolipid rheostat can be modulated with ASAH1 inhibitors to promote cell death. B. Graph showing the catalytic activity of ASAH1 protein. C. Violin plot of ASAH1 mRNA expression in different types of gliomas from TCGA database. Tukey’s Honest Significant Difference (HSD) was applied. D. Kaplan Meier plots with overall survival data from glioma patients extracted from CCGA database. Survival details and statistics are summarized in the table below. E. Western blot showing ASAH1 expression in different cell lines. β-actin was used as a loading control. E. Ceramidases mRNA expression obtained from TCGA database and RNA sequencing analysis. Friedman and Kruskal-Wallis tests, respectively, were performed (*p≤0.05; **p≤0.01; ***p≤0.001; ****p≤0.0001).

### SABRAC induces IDH^mut^-specific cytotoxicity through mitochondrial-driven apoptosis

Knowing that SABRAC was a good candidate for inducing toxicity in IDH1^mut^ cells, we performed CCK8 assays to know whether the toxicity of SABRAC was specific to IDH1^mut^ cells and to compare its efficacy with other AC inhibitors with similar structures (Fig. 2A). SABRAC and azide-SOBRAC were the more effective producing higher cytotoxicity in IDH1^mut^ (TS603 oligodendroglioma and BT142 oligoastrocytoma) compared to IDH1^wt^ (GSC923 glioblastoma) glioma cell lines (Figs. 2B, 2C and S2A). We also tested the toxicity of SABRAC and azide-SOBRAC in normal cells. The higher IC_50_ of SABRAC for non-immortalized Normal Human Astrocytes (niNHA) compared to azide-SOBRAC, identified SABRAC as a better candidate, underscoring a potentially large therapeutic window (Figs. 2C and 2D). Using western blot, we analyzed AC expression in TS603 oligodendroglioma cells under SABRAC treatment. Interestingly, we found an increase in AC only under the more toxic concentrations of the drug, suggesting that SABRAC inhibition of AC is specific and highlighting the importance of this enzyme for the survival of the cells (Fig. 2E). Additionally, we defined part of the mechanism of cell death induced by SABRAC, by treating TS603 cells with different doses of the drug (Fig. S2B). We found that both extrinsic (Fig. 2F) and intrinsic apoptotic pathways were activated. Under SABRAC treatment the Bax/Bcl-2 ratio, a cell death switch indicator (Fig. 2G), cytochrome c (Fig. 2H), cleaved-caspase 3 and cleaved-PARP1 levels (Fig. I) were increased. As ceramides can activate apoptosis through ER stress, we explored the presence of ER stress markers in SABRAC-treated conditions and found the p8 levels and the related BAD dephosphorylation enhanced (Fig. 2J). Interestingly, we did not see an increase in other typical upstream ER markers (Fig. S2C), pointing to a direct action of AC-related ceramides in mitochondrial-driven apoptosis. Similar apoptosis activation in a dose-dependent manner (Figs. S1D and S1E) was detected in NCH612 oligodendroglioma cells under SABRAC treatment.

**Fig. 2.**
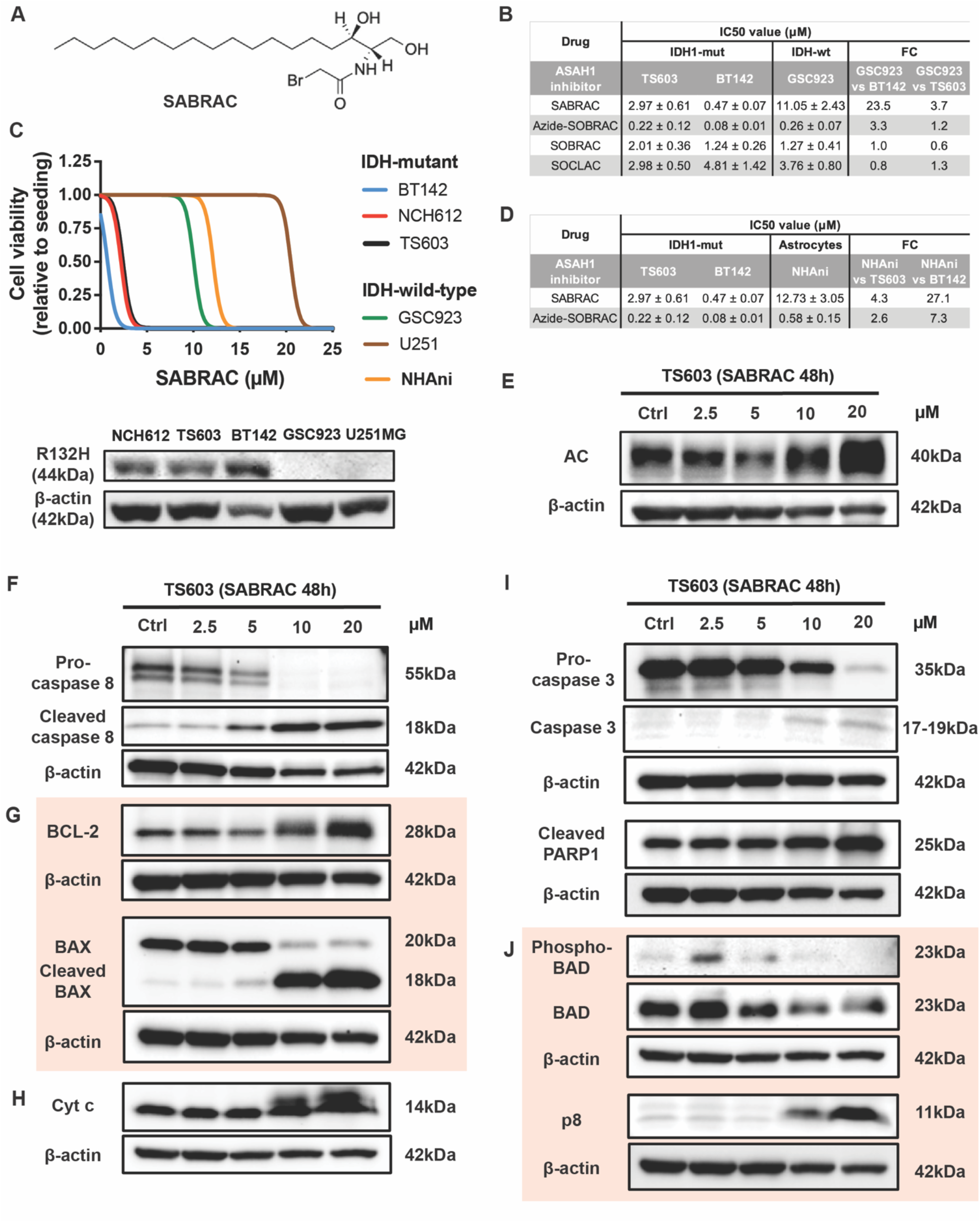
SABRAC induces IDH^mut^-specific cytotoxicity through mitochondrial driven apoptosis. A. Molecular structure of SABRAC drug. B, C (top) and D. Graph and tables compiling the IC50’s with their respective SD (for graph) and SEM (for tables) values from the ASAH1 inhibitors drug screening performed through CCK8 viability assays. The fold change columns reflect how SABRAC was the ASAH1 inhibitor more effective on producing IDH1-mutant specific cytotoxicity (A) and how SABRAC induces low toxicity to normal human astrocytes (B). Non-linear regression was applied to data. Results are shown as the mean ± SEM from at least three independent experiments. C (bottom). Western blot showing IDH1 mutation expression in different cell lines. E-J. Western blots of TS603 oligodendroglioma cells showing an increase in the expression of ASAH1 and apoptotic markers under SABRAC treatment. β-actin was used as a loading control. Images are representative of at least three independent experiments.

### SABRAC increases ceramides and their derivatives in TS603 oligodendroglioma cell lines

After confirming the mechanism of cell death of SABRAC-treated oligodendroglioma cells, we explored which specific types of ceramides were increased in these conditions activating the apoptotic process. LC/MS analysis in TS603 cells showed close to a 30-fold increase in some ceramide levels, either saturated or unsaturated. Ceramide C12 levels were most notably enhanced in a dose-dependent manner upon SABRAC addition (Fig. 3A). The increase in ceramide levels was accompanied by the activation of the salvage pathway, reflected in the enlarged amount of galactosylceramides, sulfatides, lactosylceramides, glycosphingolipids and ceramide-1-P, as well as the activation of the sphingomyelin pathway. Surprisingly, sphingosines and sphingosine-1-P levels were also found to increase under high doses of SABRAC. Another remarkable result was the activation of cholesterol pathways triggered by SABRAC treatment (Figs. 3B and 3C). To deepen in the biological consequences of AC inhibition with SABRAC, we performed RNA sequencing analysis of TS603 cells treated with non-apoptotic doses of SABRAC (2.5 μM and 5 μM). Consistent with the lipidomics results, we found increased mRNA levels of several enzymes from the salvage pathway while the mRNA levels of the main enzymes involved in the modulation of S1P levels were decreased (Fig. 3D). The effect of SABRAC in cholesterol levels seen in LC/MS results, could be partially explained with the RNA sequencing results showing impairment in cholesterol biosynthetic and transport genes (Fig. 3E).

**Fig. 3.**
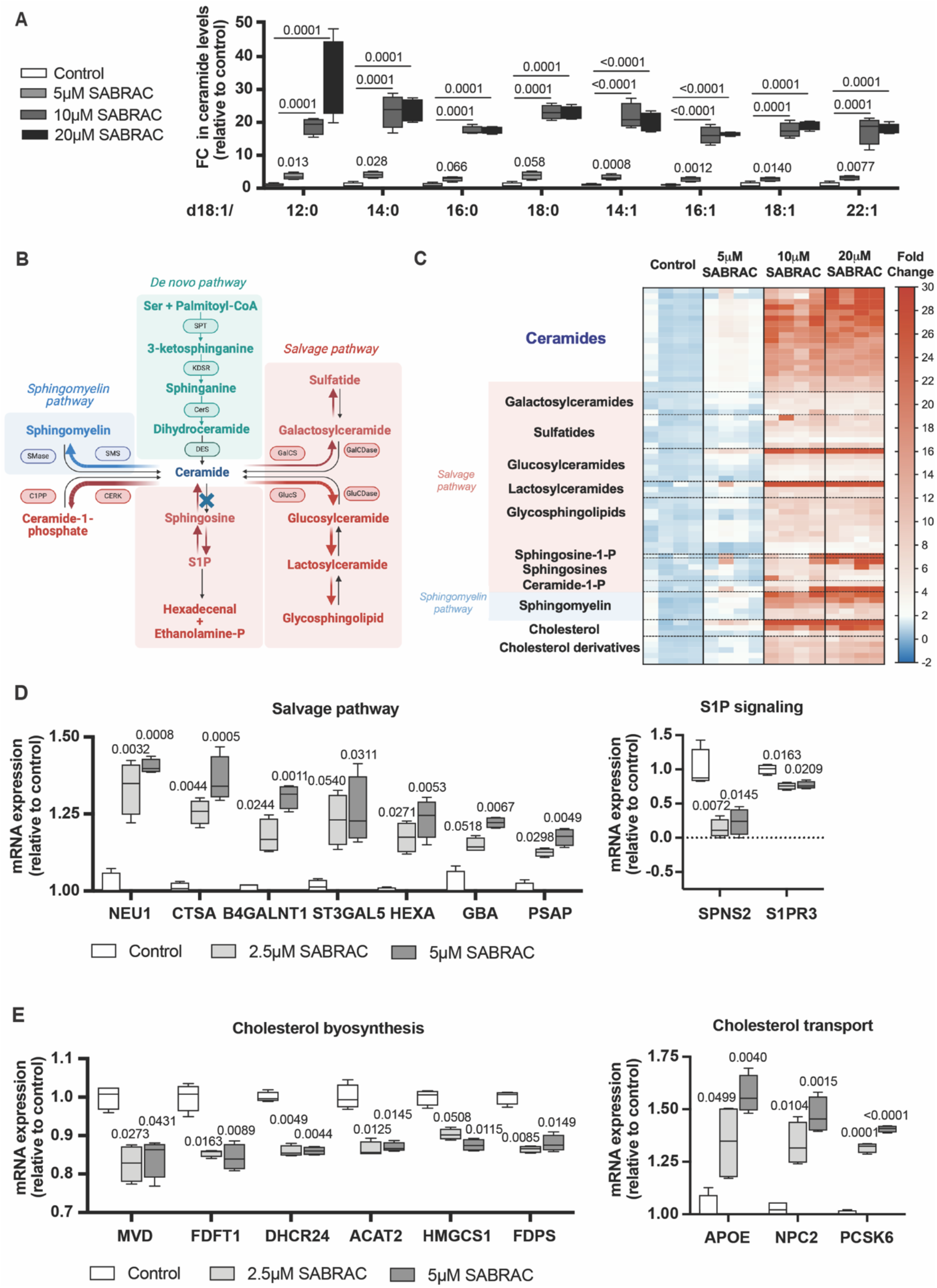
SABRAC increases ceramides and its derivatives in TS603 oligodendroglioma cell lines. LC/MS (A, B and C) and RNA sequencing (D and E) was performed in TS603 cells treated with vehicle or SABRAC drug for 48h. A. Graph bar representing ceramides increased under SABRAC treatment. Summary of four experiments is represented as mean + SD. t-test for binary comparisons and Fisher’s LSD post-hoc ANOVA for multiple group comparisons was performed. B and C. Graphical abstract and heatmap showing which lipid species and from which pathways, were found increased. D and E. Bar charts of RNA sequencing data illustrating how SABRAC modulates the expression of genes from the salvage pathway, S1P signaling, cholesterol biosynthesis and cholesterol transport. Summary of four experiments is represented as mean + SD with their corresponding p-adjusted values.

### Treatment with SABRAC prolongs survival of mice orthotopically xenografted with TS603 oligodendroglioma cells

Having demonstrated the in vitro efficacy of SABRAC in oligodendroglioma cells, we proceeded to optimize conditions and test the suitability of SABRAC for in vivo studies. We performed a preliminary repeat dose toxicity study with SABRAC under an ACUC approved animal protocol. Dosing was selected based on the work by Pearson et al.^11^, in which a drug analog from SABRAC that differs only in one atom (bromine instead of chlorine) demonstrated efficacy in *in vivo* mice studies decreasing leukemic burden by 75% ^20^. For SABRAC toxicity study, three doses of SABRAC were assessed (1mg/kg, 5mg/kg and 15mg/kg). Each animal received five intraperitoneally injections per week of SABRAC or vehicle control. Routine cage-side observations were made on all animals at least once a day throughout the study for general signs of pharmacologic and toxicologic effects, morbidity, and mortality. Mice were weighed every 3-4 days. SABRAC did not produce significant loss of weight or toxicity in the mice (Fig. S2A). 4h after the last injection mice were euthanized and plasma, liver and brain were collected. Brains were processed and analyzed by LC/MS. SABRAC was detected in a dose-dependent manner in brain tissue, indicating that SABRAC can cross the blood-brain barrier (BBB) (Fig. S2B). Once set up the conditions for the in vivo study, a mice experiment was performed for validating the in vivo efficacy of SABRAC. TS603 oligodendroglioma cells were intracranially injected and after 1 week, 15 mg/kg of SABRAC was administered intraperitoneally 5 times a week (Fig. 4A). SABRAC treated mice did not present significant weight loss respect the controls (Fig. 4B). Once completed the experiment, we found a significant improvement in survival in SABRAC treated mice with respect the control (Fig. 4C), validating the in vivo efficacy of this drug.

**Fig. 4.**
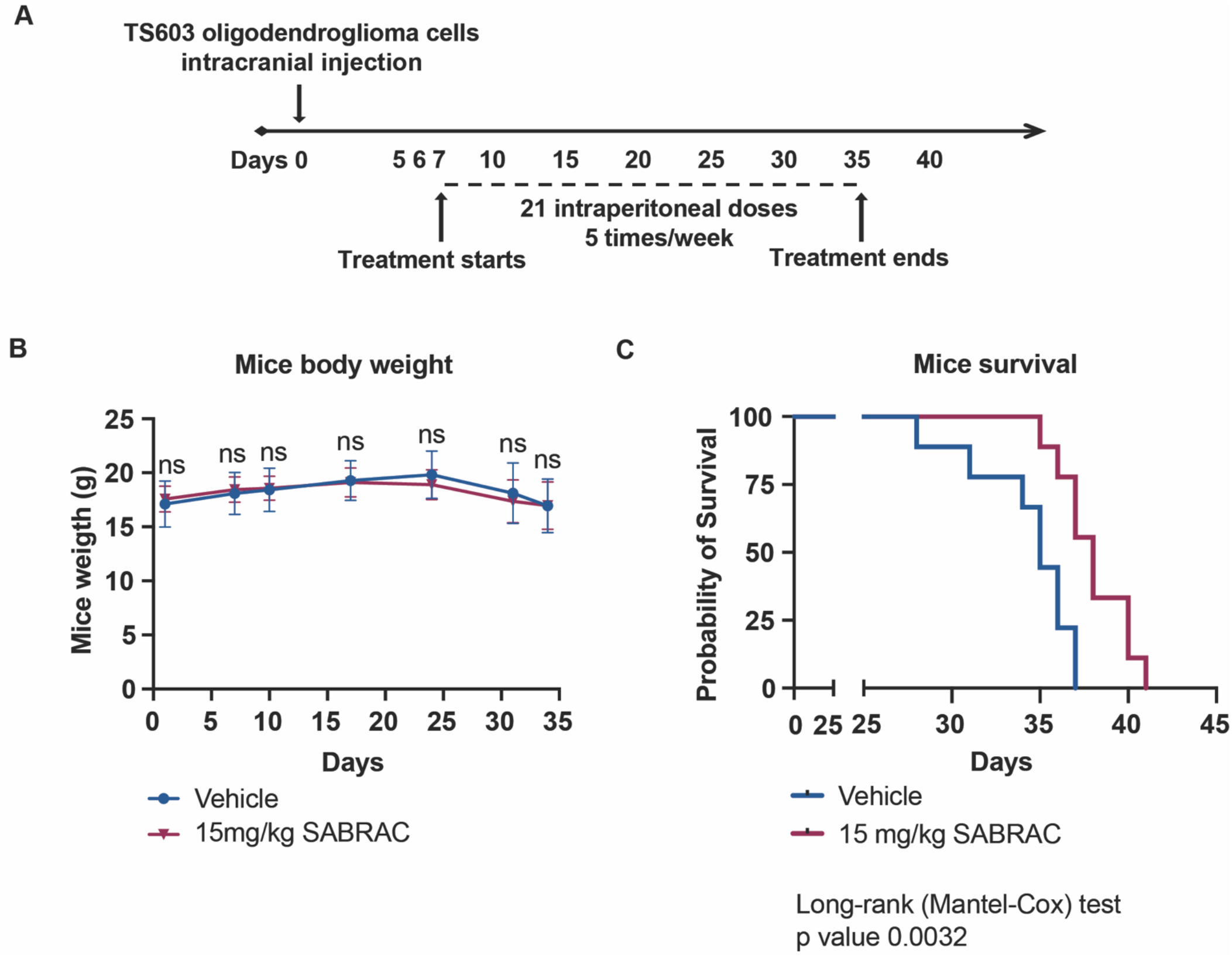
Treatment with SABRAC prolongs survival of mice orthotopically xenografted with TS603 oligodendroglioma cells. Female SCID mice (n = 9 per group) received intracranial injection of 200,000 human TS603 human oligodendroglioma cells. One week after the injection, mice were treated 5 times per week with SABRAC (15 mg/kg SABRAC in 0.5% methylcellulose and 0.2% Tween 80 in PBS) or vehicle control. A. Timeline of the in vivo experiment including the SABRAC treatment regimen. B. Graph showing that SABRAC treatment did not produce significant loss of weight in mice along the experiment. Two-way ANOVA test (ns p>0.05) was applied. C. Kaplan-Meier survival curve. Statistical significance was calculated using the log-rank Mantel-Cox test. The median survival calculated was 35 days for control mice and 38 days for SABRAC-treated mice. **p= 0.0032.

## Discussion

In this study, we demonstrate for the first time that inhibiting AC induces mitochondrial-driven apoptosis and provides a survival benefit to mice injected with IDH1^mut^ oligodendroglioma. First, we show that AC is expressed in low levels in IDH1^mut^ tumors, especially in oligodendroglioma. Using a model of U251 glioblastoma overexpressing the IDH1 mutated protein (R132), we showed that this low expression was a downstream effect of IDH1 mutation. The connection between the sphingolipid pathway and IDH1 mutation was previously identified in our group by Dowdy et al. work ^12^. There, we described how IDH1 gliomas present an impaired sphingolipid pathway that is related to increased basal levels of ceramides. To exploit this vulnerability of IDH1^mut^ gliomas, we targeted several enzymes from the sphingolipid pathway, and we identified AC inhibitors, especially SABRAC, as the most efficient drug that produced IDH1 mutant-specific cell death. Interestingly, there was a correlation between AC expression in the cell lines and sensitivity to SABRAC. SABRAC is a specific and irreversible inhibitor, therefore, it is likely that a higher expression of the enzyme will require a higher concentration of the drug to produce the same toxicity. This fact could be an advantage in terms of non-toxicity in normal human cells, as we found that normal human astrocytes express high amounts of AC. Secondly, we described the mechanism of death promoted by SABRAC in human oligodendroglioma cells. AC hydrolyzes the lysosomal membrane ceramide into free fatty acids and sphingosine. Consequently, we found that SABRAC inhibition of AC, produced an up 30-fold increase in several ceramides in TS603 oligodendroglioma cells. Levels of ceramide containing a C12 fatty acid chain were one of the more highly increased and in a dose-dependent manner upon the addition of SABRAC to cells. This data corroborates the work of Bernardo K. et al. in which they demonstrated that AC has an optimal activity for C12 ceramide and for ceramides rather than dihydroceramides ^21^. Even though ceramide’s structure is different in terms of fatty acid chain length, and their function is context-dependent (subcellular localization, and the presence of their downstream targets ^22^), several lines of evidence have established the proapoptotic role of certain ceramides ^23^. In both extrinsic and intrinsic pathways of apoptosis, several ceramides act as mediators ^24^. Cleavage of caspase-8 plays a pivotal role in the extrinsic apoptotic signaling pathway via death receptors ^25^. In our study, we found that SABRAC treatment promotes the cleavage of caspase-8 indicating its involvement in extrinsic apoptosis. Even the cleavage caspase 8 can, in turn, activate the mitochondrial apoptotic pathway ^25^, it is well-known the direct activation of ceramides to the intrinsic mitochondrial-driven pathway. Ceramides regulate apoptosis through the interaction with anti-apoptotic proteins like Bcl-xL and pro-apoptotic proteins such as Bax, and the formation of ceramide channels. Ceramide channels, contribute to the permeabilization of the mitochondrial outer membrane that leads to the release of pro-apoptotic proteins such as cytochrome c ^22^. Several markers of the intrinsic mitochondrial apoptotic pathway were found strikingly increased under SABRAC treatment in TS603 and NCH612 oligodendroglioma cells. Because ceramides can promote ER stress leading to activation of intrinsic apoptosis, we explored the expression of several ER stress markers in TS603 cells under SABRAC treatment. However, all the ER stress markers explored were found decreased, except for p8 protein and BAD phosphorylation that were increased ^26^. Consistent with the great increase in ceramides produced by SABRAC, we found a general enhancement in almost all the ceramide derivatives coming from the salvage and sphingomyelin pathways, along with an increase in the expression of several genes from the same pathways. It was surprising to find an enhancement in sphingosine and sphingosine-1-phosphate species, which however, can be explained in different ways. On the one hand, we have shown how SABRAC treatment induces an upregulation on AC expression. This is a reasonable compensatory mechanism promoted by cells for survival, especially when the activity of the protein targeted by the drug inhibitor is essential. We see that in our case, this increase in AC expression, is not able to avoid the activation of cell death, but it is likely to produce at some point an increase in sphingosines due to the breakdown of ceramides, these in turn increased because of the drug effect. On the other hand, our RNA sequencing data shows a decrease in the expression of SPNS2, a transporter that exports S1P, which could explain the significant increase of S1P inside the cells ^27^. Finally, in silico and in vitro studies have reported an AC-associated “reverse activity” that could explain increases in sphingosines until certain cellular contexts ^28^. Another surprising data obtained in this work was the increase in cholesterol in TS603 oligodendroglioma cells under SABRAC treatment. This fact was partially explained by the impairment in the expression of several genes related to the synthesis and transport of cholesterol.

In this study, for the first time, we report that SABRAC penetrates the BBB, a main challenge in the field of brain cancer therapy. Moreover, SABRAC prolongs survival in mice harboring intracranial oligodendroglioma xenograft. Despite the moderate survival benefit, this study suggests that inhibiting AC, could be considered as a potential anticancer strategy and for improving the effectiveness of other drugs that also increase ceramide levels activating apoptosis. In conclusion, we show that the SABRAC drug exerts anticancer activity against oligodendroglioma in vitro and in vivo, thereby significantly prolonging survival in orthotopic xenograft oligodendroglioma mouse models. Our results thus suggested that SABRAC represents a promising metabolically targeted treatment for patients with oligodendroglioma.

## Author Contributions

HMV, ML, TD, GK, FZ, LZ, and AL, ML have conceived, designed, performed the experiments, and analyzed the data; and conceptualized the study. HMV TD, GK, FZ, LZ, and AL has executed and validated the experiments. AL analyzed RNA seq data. MZ, WZ, HS, WZ performed the animal experiments. HMV wrote the first draft of the paper. ML supervised the study.

## Acknowledgments

We would like to thank Dr. Chun Zhang Yang (Neuro-Oncology Branch, NCI/CCR/NIH) for providing us the U251-WT, U215-R132H and U251-R132C constructs. We also thank Dr. Timothy Chan for providing us with the TS603 cell line.

## Ethical Statement

The GSC923 cell line was obtained in-house from the Neuro-Oncology Branch, NCI/CCR/NIH. BT142 was purchased from ATCC. TS603 cell line was obtained from Dr. Timothy Chan, (Cleveland Clinic). U251-WT, U215-R132H and U251-R132C constructs were obtained from Dr. ChunZhang Yang from Neuro-Oncology Branch, NCI/CCR/NIH. Intracranial orthotopic mouse models with an IDH1^*mut*^ glioma cell line were established according to an approved animal study proposal NOB-008 by the NCI-Animal Use and Care Committee (ACUC).

