## Supplemental figures for "Targeting IDH1-Mutated Oligodendroglioma with Acid Ceramidase Inhibitors"

Supplementary Information for: **Targeting IDH1-Mutated Oligodendroglioma with Acid Ceramidase Inhibitors**

[1] Helena Muley, [1] Tyrone Dowdy, [1] Faris Zaibaq, [1] George Karadimov, [1] Aiguo Li, [1] Hua Song, [1] Meili Zhang, [1] Wei Zhang, [1] Zalman Wong, [1] Lumin Zhang, [1] Adrian Lita, and [1] Mioara Larion
[1] Neuro-Oncology Branch, National Cancer Institute, Center for Cancer Research, National Institutes of Health, Bethesda, MD, USA.

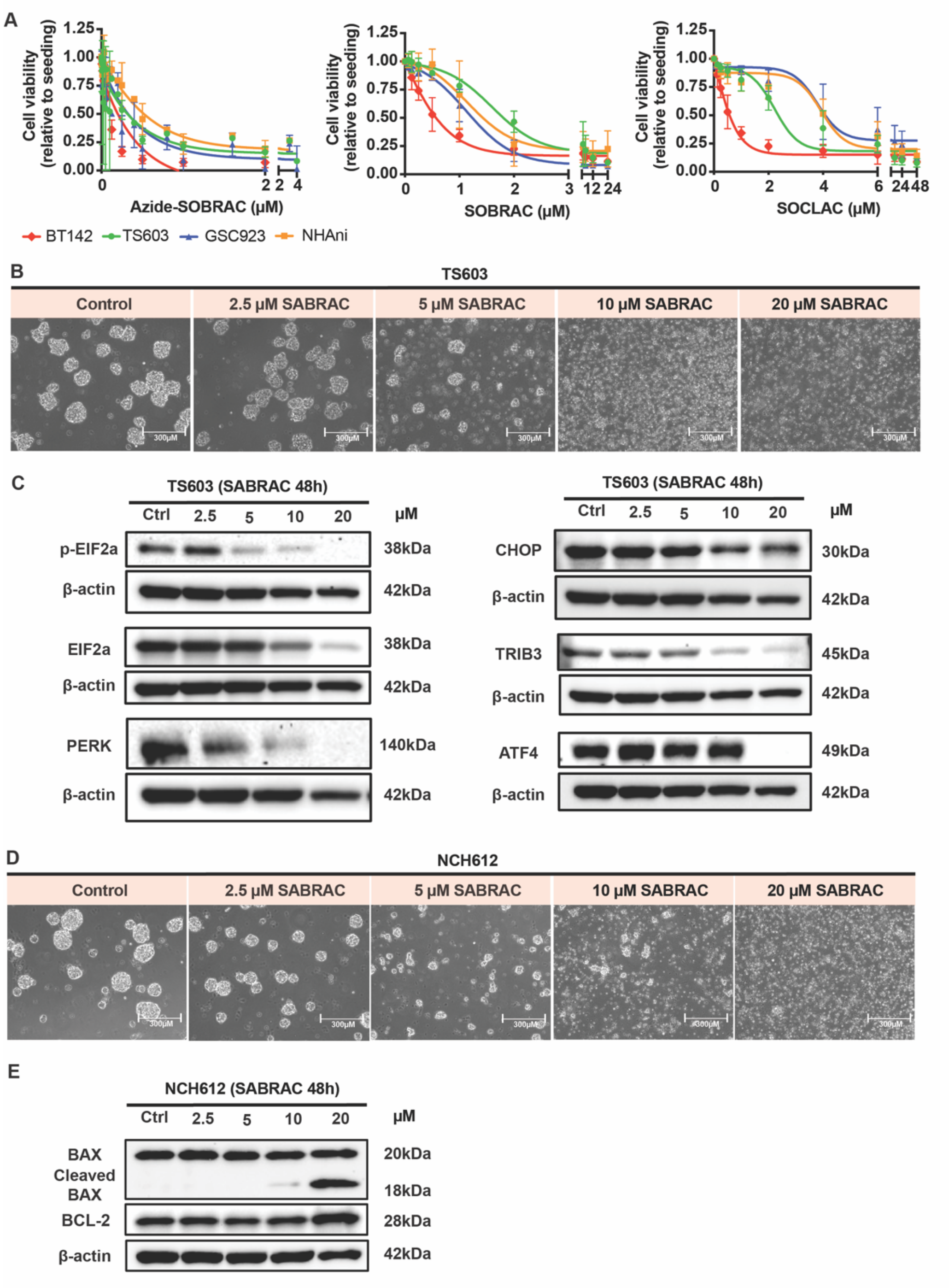

**Fig. S1. SABRAC activates apoptosis in different oligodendroglioma cell lines and does not induce ER stress in apoptotic conditions.** A. CCK8 assays for testing cytotoxicity of ASAH1 inhibitors in different cell lines. B and D. Images of TS603 and NCH612 cells after 48h under vehicle and SABRAC treatment, were acquired using an EVOS™ M5000 Imaging System microscope with a bright field objective of magnification 4×. C and E. Western blots showing different apoptotic and ER stress markers under SABRAC treatment in TS603 and NCH612 cells. β-actin was used as a loading control. Images are representative of at least three independent experiments.

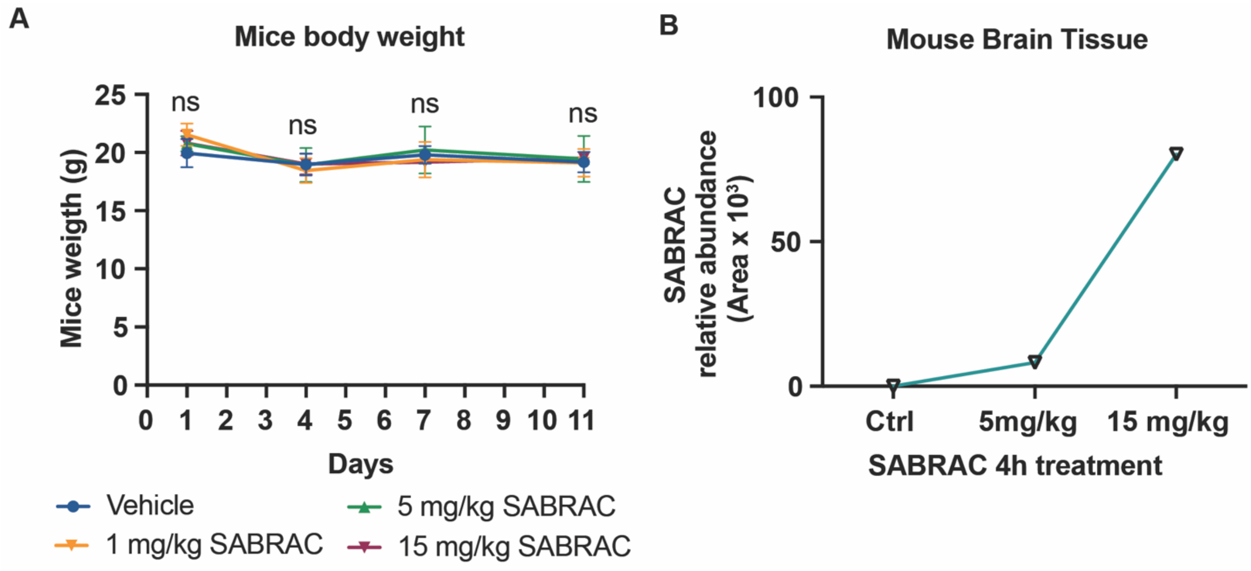


**Fig. S2. SABRAC drug passes the blood-brain barrier and does not produce a significant loss in the weight of mice.** Female SCID mice (n = 4 per group) were injected intraperitoneally five times per week with SABRAC (1, 5 or 15 mg/kg SABRAC in 0.5% methylcellulose and 0.2% Tween 80 in PBS) or vehicle control. A. Line graph showing lack of toxicity of SABRAC in mice. Two-way ANOVA test (ns $p>0.05$) was applied. B. Graph with LC/MS results showing how SABRAC was detected in brain collected 4h post intraperitoneally injection in a dose-dependent manner.
